# APEX2 and TurboID Define Unique Subcellular Proteomes

**DOI:** 10.1101/2025.09.04.673741

**Authors:** Alexandria S. Battison, Jeremy L. Balsbaugh, Jeremy C. Borniger

## Abstract

Proximity labeling has emerged as a prominent, reliable tool for obtaining local proteomes from a wide range of cell-types. Two major classes of labeling reagents, peroxidase based (APEX family), or biotin-ligase based (BioID family) have been developed in parallel. These two approaches are often used interchangeably, or chosen based on availability of reagents, however each may produce a biased proteome which should be considered during experimental design. We compared proximity labeling with TurboID or APEX2 in HEK293 cells across cytosol, nucleus, and membrane compartments. Both enzymes enriched compartment-specific proteomes, validated by GO terms, but showed distinct protein profiles. TurboID identified more membrane proteins, favoring identification of proteins associated with RNA processing and protein localization, while APEX2 enriched for proteins involved in metabolic pathways. Trypsin digestion highlighted biases from TurboID’s lysine biotinylation, which we show can be mitigated by an endoproteinase GluC digestion during sample prep, yet these differences persist to some degree. We find that TurboID suits broader proteomic studies whereas APEX2 targets specific signaling pathways. We therefore show that strategic enzyme and protease selection is critical for optimizing proximity labeling-based proteomic studies, advancing cellular proteome mapping.

## Introduction

Proximity labeling (PL) has become an indispensable tool in cell biology for mapping the molecular landscape of proteins within their native subcellular environments^1–3^. Unlike traditional co-immunoprecipitation or affinity purification approaches, which rely on capturing stable interactions and often disrupt cellular architecture, proximity labeling enables the detection of transient, weak, or spatially restricted interactions in intact living cells^1,4–6^. Proximity labeling approaches can also provide an unbiased screen of the local and transient protein environment, as they are not reliant on a lower-throughput, antibody dependent workflow in the same way as co-immunoprecipitation^6,7^. By fusing proximity labeling enzymes such as TurboID or APEX2 to a protein of interest or a targeting motif, reactive intermediates are generated that covalently tag nearby proteins with biotin^4^. These labeled proteins which represent the local, spatially constrained interacting network can then be isolated via a robust and simple streptavidin-based affinity purification and identified using mass spectrometry^8– 10^.

### Proximity Labeling Strategies

PL approaches can be broadly categorized into two families: Biotin-ligase based approaches and peroxidase-based approaches, discussed in detail below. Biotin-ligase based approaches include BioID, BioID2, and TurboID, with peroxidase-based approaches including APEX, APEX2 and other horse-radish peroxidase dependent labeling strategies.

### Biotin-Ligase based approaches

BioID, introduced in 2012, is derived from a mutated form of the *Escherichia coli* biotin ligase BirA, which promiscuously biotinylates lysine residues on proteins within a ∼10 nm radius when supplied with exogenous biotin^11,12^. Early applications of BioID included elucidating previously challenging subcellular regions, such as the centromere-cilium interface, or transient protein-protein interactions in yeast^13,14^. However, BioID is not without limitations, most notably the long incubation time needed for labeling (on the order of hours) prevents precise temporal control of the labeling radius^15^. To address these slow labeling kinetics, TurboID was developed^16^. TurboID includes a highly active *E. coli* BirA variant with significantly faster biotinylation kinetics on the order of minutes, capable of achieving labeling significantly faster compared to the hours needed for BioID and BioID2^15,17,18^. TurboID represents a significant advancement in proximity-dependent biotinylation techniques for studying protein-protein interactions and subcellular proteomes in living cells^19,20^ This faster labeling rate allows for more precise control over the labeling radius and increased temporal precision. These successive innovations have made TurboID a powerful tool for mapping dynamic protein interactions and subcellular environments with high temporal and spatial resolution^21^. TurboID has been pivotal in revealing low-abundant protein complexes in *Arabadopsis*, as well as subcellular compartments in neurons and microglia and subsequent dynamic changes in immune responses within cells following lipopolysaccharide (LPS) treatment^22,23^. Together these studies highlight the broad utility of biotin-ligase based approaches, particularly *in vivo*, for capturing subcellular compartments that would otherwise be challenging to isolate by fractionation methods alone, or whose interactions are too transient to be captured by immunoprecipitation.

### Peroxidase-based approaches

During the same time that BioID was developed, a peroxidase-based approach for proximity labeling was introduced. The peroxidase family of proximity labeling enzymes includes APEX and APEX2, which make use of an engineered ascorbate peroxidase and exogenously supplied biotin-phenol to label tyrosine residues^24,25^. Upon addition of hydrogen peroxide (H_2_O_2_) and the biotin-phenol substrate, APEX2 catalyzes the one-electron oxidation of biotin-phenol to a short-lived biotin-phenol radical^26,27^. This highly reactive species covalently labels the electron-rich amino acid tyrosine on proteins within a ∼20 nm radius of the enzyme^6,8^. Biotinylated proteins are then enriched via streptavidin affinity purification and identified by mass spectrometry. Similar to TurboID, APEX2 labeling occurs within seconds, making it well-suited for capturing dynamic interactions and subcellular proteomes with high temporal resolution^28^. The fast labeling made possible by APEX has been instrumental in tracking G-protein coupled receptor (GPCR) signaling dynamics, which occur within seconds and are challenging to capture by any other approach^29^.

Both biotin-ligase and peroxidase-based methods have opened the door to studying spatially defined proteomes from subcellular compartments which may be challenging to impossible to isolate by organelle fragmentation methods alone. As a result, many researchers have successfully employed one of these two strategies to label a local proteome within a specific subcellular compartment^7,18,30,31^. Hung et al. (2016) used APEX2 targeted to the mitochondrial matrix to uncover differences in mitochondrial proteomes across tissues, revealing novel tissue-specific metabolic pathways^24^. Branon et al. (2018) employed TurboID to systematically define the proteomes of multiple subcellular compartments in mammalian cells, providing a comprehensive atlas of spatial protein organization under physiological conditions^32^. Despite the wide adoption of these powerful techniques, direct head-to-head comparisons of these two PL reagents and the proteomes which result has not been fully established. Here we sought to drive expression of either APEX2 or TurboID across three commonly labeled subcellular compartments (cytoplasm, nucleus, and membrane) and compare the proteomes that result from each PL enzyme. This allows us to determine whether any biases exist in which proteins and biological components they are capable of capturing.

## Results

APEX2 or TurboID were directed to specific cellular compartments by transfecting HEK293 cells with plasmids directing PL to either the cytosol (Nuclear Export Sequence, NES), nucleus (Nuclear Localization Sequence, NLS), or membrane (CAAX/Gp41 motif) (Fig 1A,B). Immunohistochemistry was performed to first validate both plasmid transfection and correct subcellular localization (Fig. 1C). Following successful validation, transfected HEK cells were biotinylated according to the appropriate protocol for either APEX2 or TurboID (see methods for details). Briefly, APEX2 transfected cells were biotinylated through the addition of a biotin-phenol and hydrogen peroxide for 1 minute before quenching the labeling reaction with sodium ascorbate. TurboID transfected cells were biotinylated by adding biotin and with the addition of ATP for 1 minute, samples were biotinylated (Fig 1D). Successful biotinylation was validated using FITC-streptavidin immunohistochemistry. Samples were lysed and homogenized and pulled down on streptavidin beads to enrich for biotinylated proteins, which were identified using untargeted liquid chromatography-tandem mass spectrometry (LC-MS/MS).

**Figure 1.**
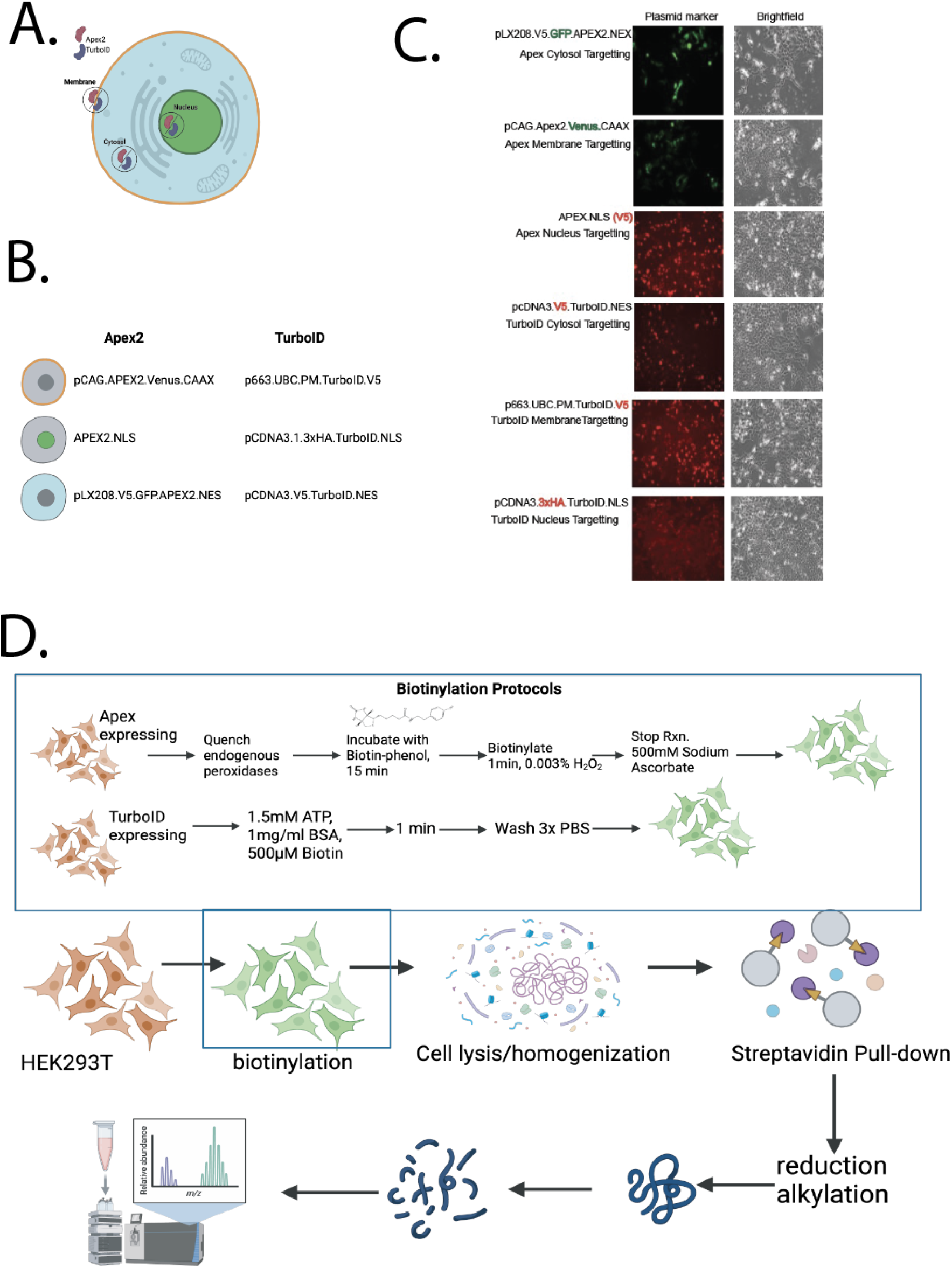
A) Schematic of cellular compartments targeted by APEX2 and TurboID B) plasmid systems used to target subcellular compartments. C) Immunohistochemistry validation of plasmid transfection using either GFP tag within plasmid, or V5 or HA tag (red). D) Schematic of protocol used for biotinylating using either APEX2 or TurboID. *Schematics made by the authors using BioRender*.

### Validation of compartment enrichment

We confirmed successful enrichment for each subcellular compartment targeted by each plasmid. Comparing the list of identified proteins across APEX2 or TurboID, we note that while there are proteins shared between the two PL approaches, there are many proteins uniquely identified in each, suggesting that different proteins are being enriched when one PL labeling approach is used over the other (Fig 2A). Gene Ontology (GO)^33,34^ term annotation and enrichment analysis demonstrates that both APEX2 and TurboID enriched for a proteome that correctly reflects the subcellular compartment we were intending to label (Fig 2B-D). We confirm cytoplasm-associated GO cellular compartment terms (cytosol, cytosolic ribosome, cytosolic region) as well as terms associated with processes occurring in the cytoplasm, such as eukaryotic translation using both methods (Fig 2B). When targeting APEX and TurboID to the membrane, we see membrane associated gene ontology terms including plasma membrane, mitochondrial outer membrane, and secretory granule membrane. We do note that many organelle membranes are captured as well including lysosomes, mitochondria, and the cytoplasmic periphery of the nuclear pore (Fig 2C). Finally, we see enrichment for nuclear GO terms when APEX2 or TurboID are driven to the nucleus via a NLS sequence, including nuclear lumen, nucleus, nuclear speck, and snRNP complexes (Fig 2D). Collectively our initial gene ontology profiling confirms that we successfully targeted the subcellular compartment we were intending to using both PL enzymes.

**Figure 2.**
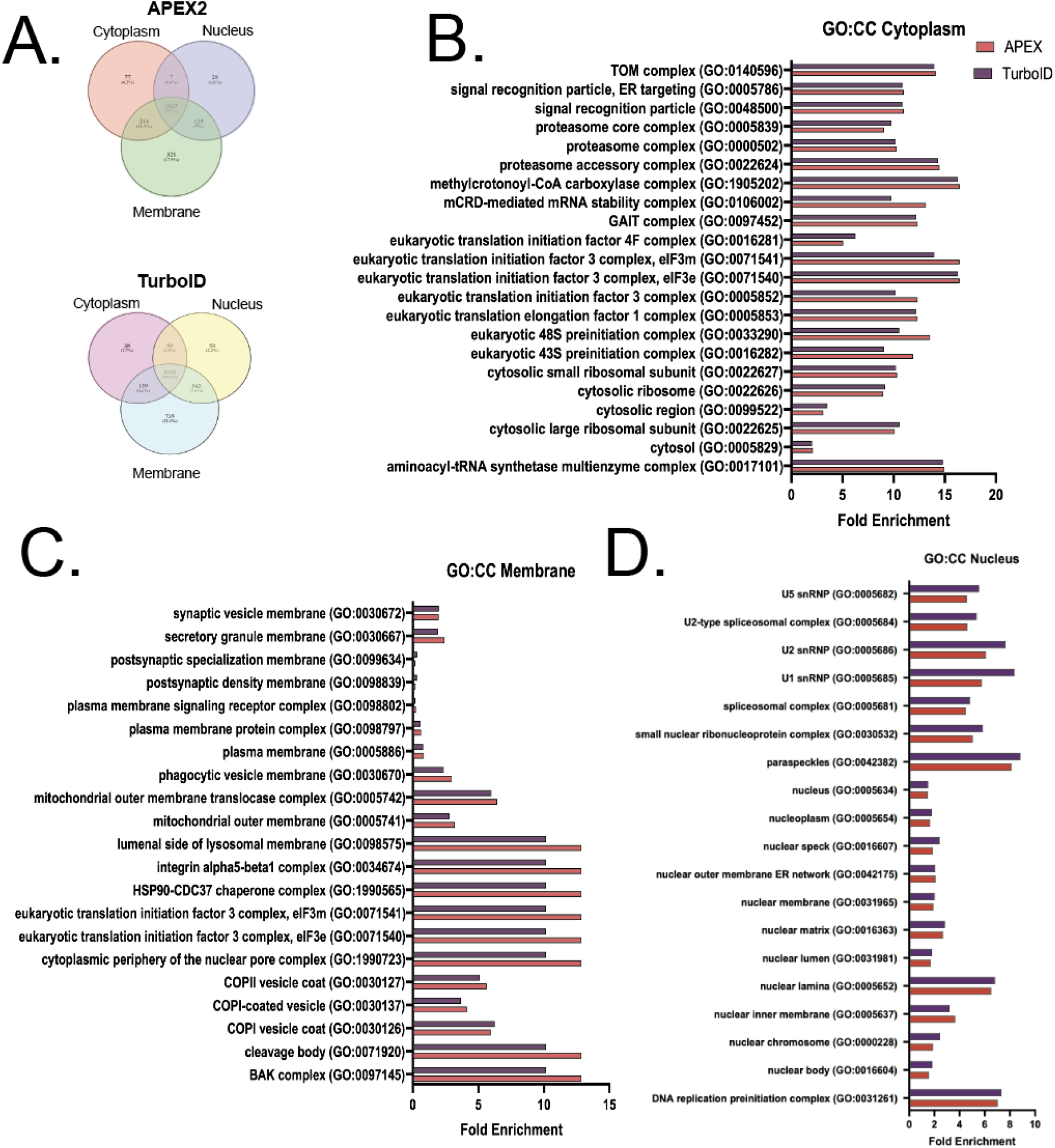
A) Venn Diagrams of total proteins identified using each approach by cellular compartment. B) Gene ontology cellular component (GOCC) terms shared between APEX2 and TurboID and the respective fold enrichment values C). GOCC fold enrichments for labeling the membrane and D) GOCC fold enrichments for labeling the nucleus with APEX2 and TurboID

### APEX2 and TurboID create local proteomes

After establishing that our constructs are targeting the intended compartments of the cell, we sought to validate that a local proteome was in fact being targeted. Across all samples analyzed, the total number of proteins identified ranges from 371 to 1266, and all intensities fall within the same range (Fig 3A, B). We next performed pairwise comparisons comparing each compartment and proximity labeling approach and confirm that, relative to a control whole cell lysate, a compartment specific enrichment was observed for each combination of compartment and proximity labeling method (Fig 3C). We next wanted to compare APEX2 to TurboID within a compartment-specific enrichment, and observed that directing APEX2 and TurboID to the membrane produced the most distinct differential enrichment of proteins (Fig 3D).

**Figure 3.**
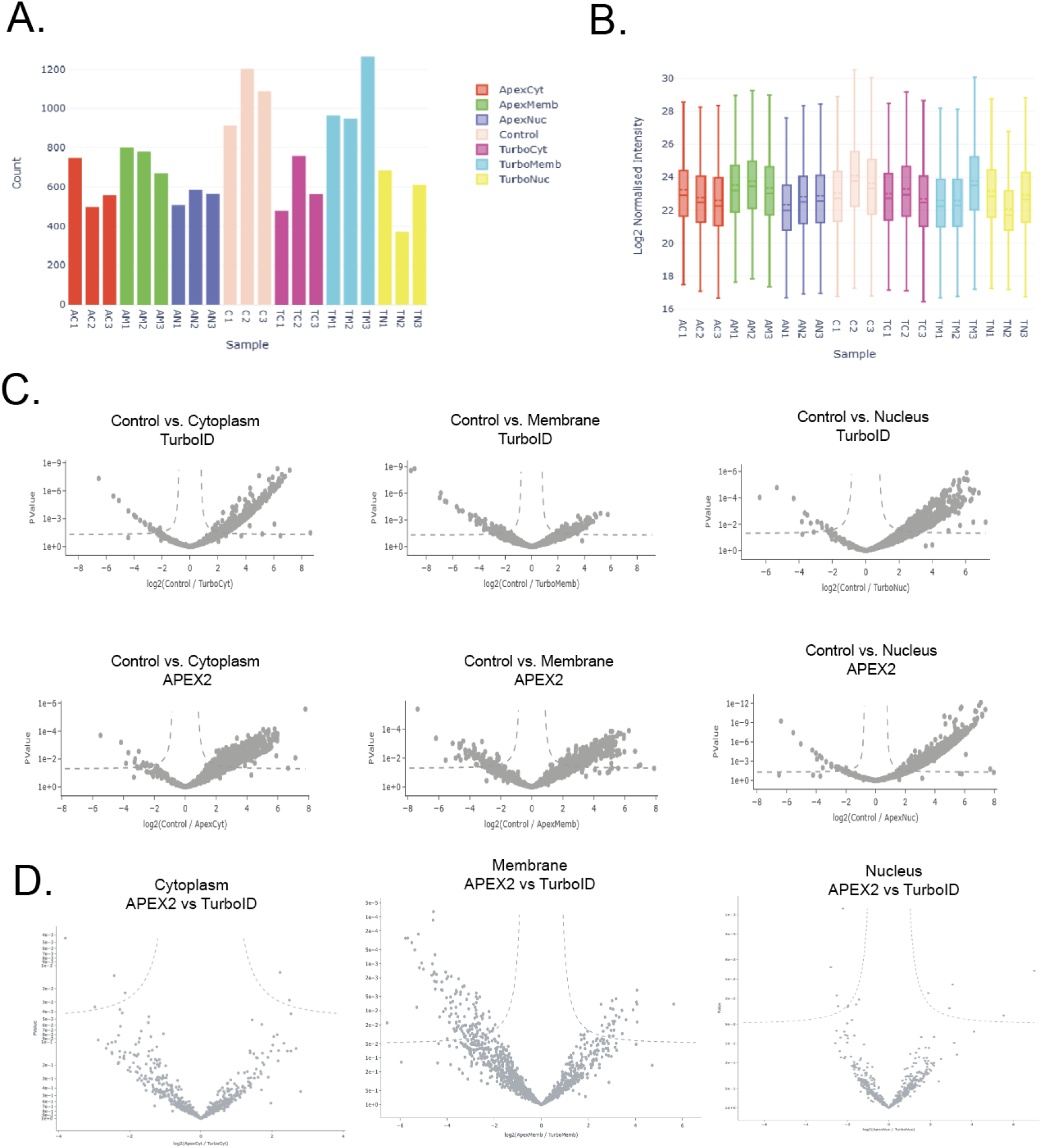
A). Total protein count for all samples and cell-lysate control B). Log2 normalized intensities for all samples and controls C). Pairwise comparison of cytoplasm, membrane, or nuclear enrichment for TurboID (top) or APEX2 (bottom). D). Volcano plot of cytoplasm, membrane, or nuclear enrichment when compared between APEX2 and TurboID.

### Biased proteomes generated from different PL reagents

Broadly, we demonstrate that the cellular compartment (GO:CC) terms are nearly identical for either APEX2 or TurboID when being directed towards the same cellular compartment (Fig 2A-D). However, we do note that there is not a complete overlap between the two and some unique cellular compartments are found (Fig 4A). APEX2 and TurboID when directed to the cytoplasm share 73% of the gene ontology terms, however each have approximately 50 terms which are unique to that enrichment method. Additionally, APEX2 and TurboID share approximately 70% of GO:CC terms when expressed on membranes, however APEX2 only has 19 unique GO:CC terms whereas TurboID has 140, suggesting that there is a difference in enriched membrane proteomes. When expressed in the nucleus, APEX2 and TurboID have more similar GO:CC enrichment, however 34 unique terms are found for APEX2 and 66 for TurboID, again suggesting that there are nuanced differences in the proteomes that are obtained with each approach.

**Figure 4.**
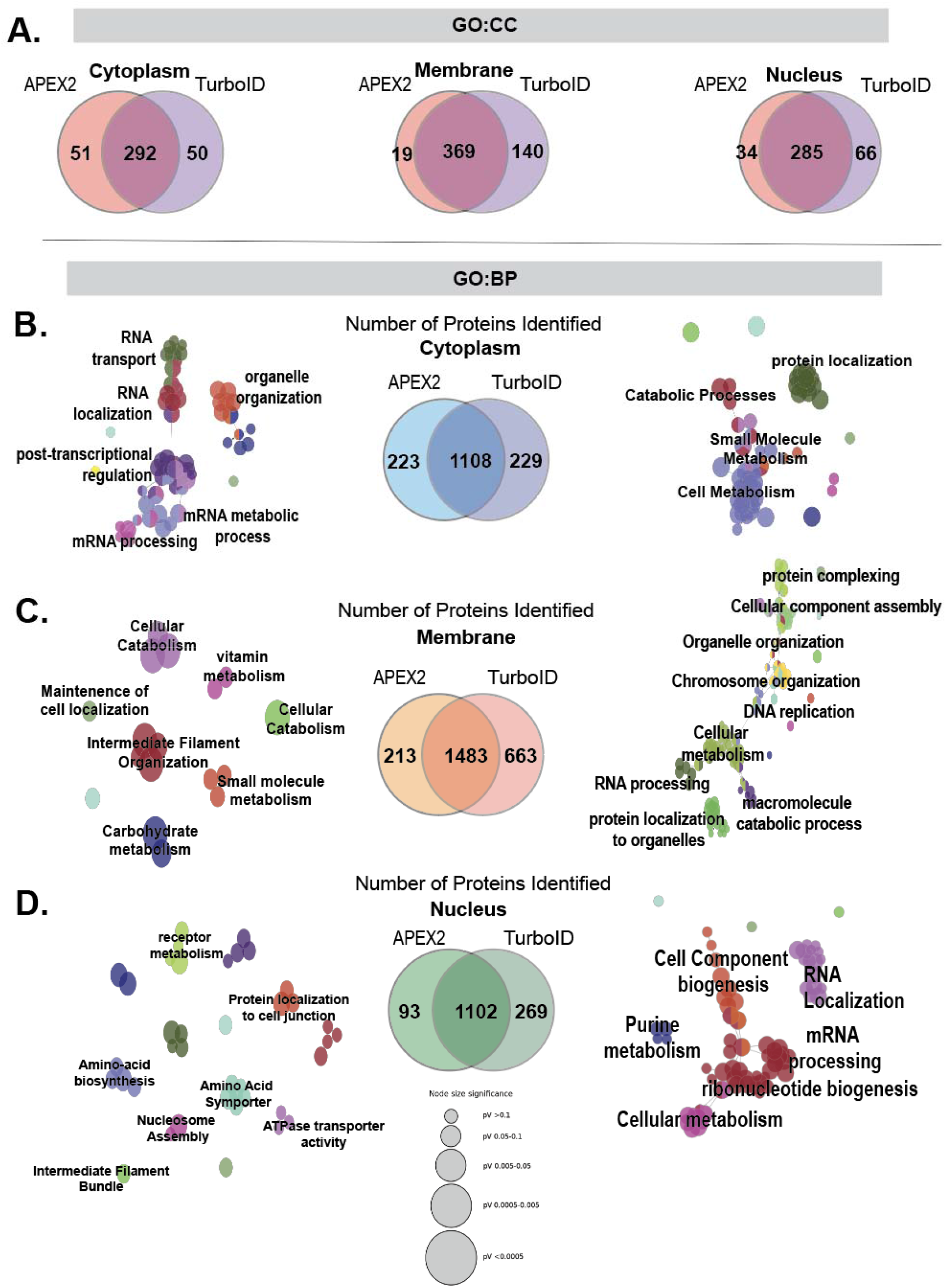
A) Venn-Diagrams of cellular component (GO:CC) terms identified for cytoplasm, membrane, and nuclear localization when using either APEX2 or TurboID. B). Biological Process (GO:BP) networks (cytoscape) and total number of proteins identified for cytoplasm C) Membrane and D) nuclear subcellular compartments. Bubble size indicates p-value.

To investigate what differences exist between these two proteomes, we quantified the total number of proteins identified for APEX2 or TurboID for each subcellular compartment. As in our GO:CC enrichment results, we observe that most proteins are shared but depending on labeling approach and compartment, between 93-663 are uniquely identified (Fig 4B-D). This again suggests that these two PL approaches do not enrich identical local proteomes. We used GO biological process (GO:BP) to delineate the functional classes of unique proteins identified using each labeling method. Across GO:CC, total protein, and GO:BP analyses, APEX2 and TurboID are most comparable when expressed in the cytosol. There are 223 (APEX2) and 229 (TurboID) proteins which are uniquely identified using one PL approach over the other. These unique proteins reflect differences in biological process enrichment, with APEX2 capturing proteins related to RNA localization and transport and mRNA processing, whereas TurboID mainly captures proteins associated with cell metabolism (Fig 4B).

The biggest difference between APEX2 and TurboID appears to occur when capturing the local proteome around a membrane. There is the largest discrepancy in the number of proteins captured, with APEX2 capturing 1696 unique proteins compared to 2146 by TurboID (Fig 4C). TurboID captures a host of biological processes spanning from RNA processing to protein complexing and protein localization, reflecting perhaps an enhanced ability to capture membrane bound proteins across numerous cellular membranes. APEX2 in contrast shows enrichment for biological pathways related to small molecule metabolism or intermediate filament organization, but no membrane specific signaling pathways.

Finally, nuclear enrichment shows a discrepancy in the number of proteins identified as well, with 93 proteins being uniquely identified by APEX2 and 269 uniquely identified by TurboID (Fig 4D). We observe that APEX2 shows a modest enrichment for biological processes such as nucleosome assembly, intermediate filament bundle assembly, and amino acid biosynthesis, whereas TurboID shows a much more substantial increase in terms like mRNA processing, RNA localization, and Purine metabolism.

### APEX2 and TurboID labeling followed by enrichment leads to distinct quantitative profiles for local proteomes

We compared proteins that were identified by APEX2 and TurboID using a tryptic digest and found that broadly, there was little correlation between the protein intensity and identification between the two PL approaches, regardless of which compartment was biotinylated. This is not driven by amino acid frequency, as amino acid frequency from all peptides identified is distributed in a similar manner across proximity labeling approach and protease (Supplementary Figure 1). Given that trypsin cuts C-terminal to lysine and arginine residues (unless followed by a proline, Fig 5A), and that biotin-ligase based approaches such as TurboID biotinylate lysine residues, we suspected multiple missed cleavages post-trypsinization may lead to fewer modified peptide identifications. Therefore, we tested a non-lysine targeting alternate protease to increase protein and protein IDs and intensities. We proteolyzed a new set of samples with the serine protease endoproteinase GluC (NEB), which when buffered in ammonium bicarbonate cuts C-terminal to glutamic acid, a residue which is not biotinylated by either APEX2 or TurboID, (Fig 5B). We see stronger correlations between APEX2 and TurboID identified proteins when using GluC compared to trypsin (Fig 5C,D). There are several potential explanations for this: (1) unique quantifiable peptides are created with GluC which are not created with trypsin that are not susceptible to poor cleavage efficiencies resulting from biotinylation; (2) the GluC digestion may cleave in a manner that is more reproducible and less dependent on protein structure or buffer pH; (3) or conversely trypsin may over digest some primary sequence regions that GluC preserves. Most crucially, the use of either protease does not produce a one-to-one mapping of the proteomes obtained by either method, suggesting that the bias persists independent of sample prep and is intrinsic to the labeling reagents themselves.

**Figure 5.**
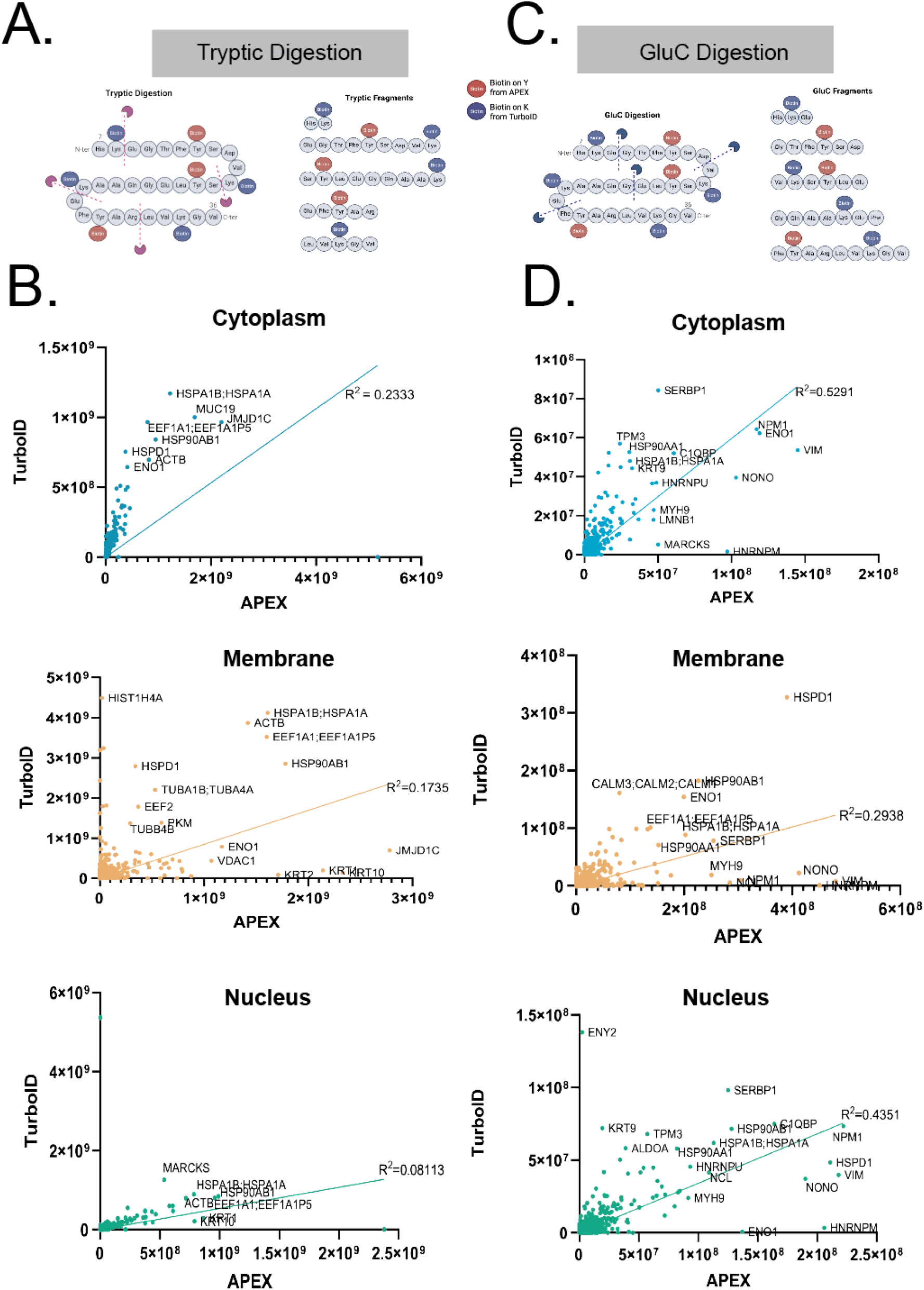
A) Schematic of location of biotinylated residues using either APEX (red circles) or TurboID (blue circles) on an example peptide. Tryptic cut sites shown as purple icon and dashed lines. B). XY plots of protein intensities after tryptic digestion for APEX2 vs TurboID. R^2 = 0.2333 for cytoplasm, 0.1735 for membrane enrichment and 0.08113 for nucleus. C) schematic of endoproteinase GluC digestion on an example peptide. GluC cut sites are shown by the blue icon and dotted line D) XY plots for protein intensities after GluC digestion for APEX2 vs TurboID. R^2 = 0.5291 for cytoplasm enrichment, 0.2938 for membrane, and 0.4351 for nuclear.

## Conclusions

Proximity labeling (PL) techniques, such as TurboID and APEX2, have revolutionized the study of protein interactions and subcellular proteomes by enabling the capture of transient and spatially restricted molecular interactions in living cells^35–37^. Our study provides a direct comparison of these two prominent PL enzymes across three subcellular compartments, cytosol, nucleus, and membrane, in HEK293 cells, revealing both shared and distinct proteomic profiles. While both TurboID and APEX2 successfully enriched for compartment-specific proteomes, as validated by gene ontology (GO) cellular compartment terms, significant differences in the proteins identified and their associated biological processes highlight inherent biases in each method. These findings underscore the need for careful consideration of enzyme choice based on experimental goals.

The observed proteomic differences between TurboID and APEX2 likely stem from their distinct biochemical mechanisms. TurboID, a biotin ligase, promiscuously biotinylates lysine residues within a ∼10 nm radius, with rapid kinetics enabling labeling in minutes. In contrast, APEX2, an ascorbate peroxidase, generates short-lived biotin-phenol radicals that primarily label tyrosine residues within a ∼20 nm radius in seconds. These differences in labeling chemistry and spatial reach likely contribute to the unique protein subsets identified by each enzyme. For instance, TurboID enriched more membrane-associated proteins (2146 unique proteins vs. 1696 for APEX2), potentially due to its ability to label lysine-rich proteins across diverse membrane environments, as evidenced by broader GO biological process terms spanning RNA processing to protein localization. Conversely, APEX2 showed enrichment for processes like small molecule metabolism, suggesting a bias toward specific protein classes or microenvironments.

Analysis using alternative proteases (trypsin vs. GluC) further confirms these biases and demonstrate that they are independent of sample prep approach. Trypsin, which cleaves C-terminal to lysine and arginine, showed poor correlation in protein intensities between TurboID and APEX2, likely because biotinylation of lysine residues from TurboID interferes with efficient tryptic cleavage. Switching to GluC, which targets glutamic acid and is less affected by biotinylation improved correlation but did not eliminate differences, indicating that biases are intrinsic to the labeling mechanisms rather than solely a result of sample preparation. This suggests that researchers should consider protease choice alongside PL enzyme selection to optimize proteomic coverage.

These findings have important implications for PL-based studies. While TurboID and APEX2 both offer high spatial and temporal resolution compared to traditional methods like co-immunoprecipitation, their proteomic outputs are not interchangeable. Researchers targeting membrane or nuclear proteomes may prefer TurboID for its broader coverage, whereas APEX2 may be better suited for specific metabolic pathways or tyrosine-rich protein environments. Future studies could explore hybrid approaches or complementary use of both enzymes to achieve more comprehensive proteomic maps. Additionally, further investigation into the structural and chemical properties of labeled proteins could clarify the molecular basis of these biases, enhancing the precision of PL techniques in dissecting cellular proteomes.

## Methods

### Plasmids

Targeting of proximity labeling reagents to the nucleus was accomplished through transfecting with Lipofectamine 2000 (Thermo) either a APEX.NLS (Addgene Cat. No 124617) or pcDNA3.1.3xHA.TurboID.NLS (Addgene Cat. No 107171) plasmid. Membrane targeting was accomplished using pCAG.APEX2.Venus.CAAX (Addgene Cat. No 168511) or p663.UBC.PM.TurboID.V5 (Addgene Cat. No 214455) plasmid. Cytoplasmic targeting was accomplished with a pLX208.V5.GFP.APEX.NES (Addgene Cat. No 202018) or pCDNA3.V5.TurboID.NES (Addgene Cat. No 107169) plasmid.

### Cell Culture

HEK293 cells (ATCC) were cultured in DMEM (Gibco) with 10% Fetal Bovine Serum (FBS, Gibco) and 1% PenStrep (Gibco). Cells were transfected using lipofectamine 2000 (Thermo), and FBS was dropped to 1% in transfection media to reduce the amount of free-biotin present. 10μg of plasmid was transfected per dish. Cells were maintained at 37C with 5% CO2. 48 hours after transfection cells were harvested and fixed with 4% paraformaldehyde.

### Biotinylation

TurboID biotinylation was performed by incubating cells with 1.5mM ATP, 1mg/mL BSA, and 500μM biotin for 1 minute before washing 3 times with phosphate-buffered saline (PBS), pH 7.4.

APEX2 biotinylation was performed by quenching endogenous peroxidases by incubation with 0.5% H_2_O_2_. After wash out of H_2_O_2_, samples were incubated with biotin-phenol (biotin-tyramide, Invitrogen, Cat No B40951) for 15 minutes before catalyzing biotinylation through the addition of 0.003% H_2_O_2_ for 1 minute. Biotinylation was quenched using 500mM sodium acetate.

### Immunohistochemistry validation

Successful biotinylation and plasmid transfection was validated using immunohistochemistry. To validate that plasmid transfection was successful, either GFP from the plasmid was visualized or staining was performed with either an anti-V5 (Invitrogen, Cat No R960-25) or anti-HA antibody (Invitrogen Cat No. 26183). Successful biotinylation was confirmed using FITC-Streptavidin (Invitrogen, SA10002) or Streptavidin-568 (Invitrogen Cat No. S11226).

### Sample lysis and pull-down

Following confirmation of biotinylation and transfection, samples were prepared for mass spectrometry. Sample lysis buffer contained 1% w/v N-octylglucoside (Thermo Cat No 28310), 1% w/v CHAPS (Thermo Cat No. 28300) 0.5% w/v sodium deoxycholate (Therno Cat No 30970), 50mM Tris, 1M NaCl (Thermo AM9760G) and 1M MgCl_2_ (Themo Cat No. AM9530G). Samples were lysed for 2 hours then spun at 16,500xg for 10 minutes. Supernatant was then added to cleaned and washed streptavidin-M280 Dynabeads (Thermo Fisher, Cat No 11205D) and incubated with beads for 2 hours. Samples were washed three times with 1x PBS, once with 1M NaCl, then 3 more times with PBS before being resuspended in 150μl 50mM ammonium bicarbonate, pH 8.5 (AmBic, Thermo Cat No. 399212500)

### Mass spectrometry sample preparation

Samples were prepared as previously described^38,39^. Samples were reduced by adding 5μl of 250mM dithiothreitol (Thermo Cat No. 20290) for 1 hour, then alkylated by adding 12.5μl of 250mM iodoacetamide (Thermo Cat No.122270250) for 45 minutes in the dark. Samples were proteolyzed on beads using either trypsin (Promega Cat No 90057, 1:20 E:S ratio) or GluC (NEB P8100S, 1:20 E:S) overnight at 37^°^C. Peptides were then acidified and desalted using Phenomenex SPE cartridges (Cat No 8B-S100-AAK) according to manufacturers specifications. After desalting, peptides were dried to completion, resuspended in 0.1% formic acid in water, and quantified on a UV-VIS Spectrophotometer using A280/A260 (1 Abs = 1 mg/ml). Loading amounts were normalized across all samples based on A280 readings.

### LC-MS/MS

Peptide samples were subjected to mass analysis using a Thermo Scientific Ultimate 3000 RSLCnano ultra-high performance liquid chromatography (UPLC) system coupled to a high-resolution Thermo Scientific Orbitrap Eclipse Tribrid mass spectrometer, using methods previously established^38,40^. Each sample was injected onto a nanoEase M/Z Peptide BEH C18 column (1.7μm, 75μm x 150mm, Waters Corporation) and separately by reverse-phase UPLC using a gradient of 4-30% solvent B (0.1% formic acid in acetonitrile) over a 90-minute gradient at 300 nl/min. Peptides were eluted directly onto the Eclipse Tribrid using positive mode ESI. MS scan acquisition parameters included resolution 60,000, AGC target 1e6, maximum injection time (MIT) of 60ms, and mass range of 300-1800 m/z. Data-dependent MS/MS spectra were acquired in the linear ion trap (IT) using collision-induced dissociation (CID) and the following parameters: Dynamic exclusion “on” with a window of 30 sec, charge state inclusion of +2 to +8 charge states only, exclude after n times set to 1, intensity threshold of 5.0e3, cycle time of 3 sec, isolation window of 1.6 m/z, 35% CID collision energy and 10 ms CID activation time, “Rapid” IT scan rate, “Normal” mass range, “Standard” AGC target, and “Dynamic” MIT. For ETD fragmentation, spray voltage in positive ion mode was 2200V. Cycle time of 3 sec, included charge states 3-19, included undetermined charge state = false. Dynamic exclusion after 1 times, exclusion duration of 30 sec, mass tolerance = ppm, mass tolerance low = 10, high = 10. The m/z isolation window = 1.6 m/z, activation type = ETD, use calibrated charge dependent ETD parameters = true. Detector type = orbitrap, orbitrap resolution is 60K with a MIT 118 msec and AGC 50000.

### Data Analysis

Peptides were identified and quantified for all experiments with the Andromeda search engine embedded in MaxQuant v1.6.0.1.^41^ Raw data were searched against the UniProt *Homo sapiens* reference proteome (UP000005640, Accessed 12/12/2024) plus the MaxQuant contaminants database. Oxidation(M), acetylation(protein N-term) were set as variable modifications. Carbamidomethylation(C) was set as a fixed modification. Protease specificity was set to trypsin/P or GluC with a maximum of 2 missed cleavages. LFQ quantification was enabled and minimum number of amino acids/peptide set to 5. All results were filtered to a 1% false discovery rate at both the peptide and protein levels based on a target/decoy method; all other parameters were left as default values.

The raw LC-MS/MS data for identifiying biotin modifications was searched using MetaMorpheus v1.1.2 with G-PTM search^42,43^. The following search settings were used: protease = trypsin or GluC (per each experimental condition); search for truncated proteins and proteolysis products = False; maximum missed cleavages = 2; minimum peptide length = 7; maximum peptide length = unspecified; initiator methionine behavior = Variable; fixed modifications = Carbamidomethyl on C, Carbamidomethyl on U; variable modifications = Oxidation on M; Biotinylation on K, Biotin on K; G-PTM-D modifications count = 2; precursor mass tolerance(s) = {±5.0000 PPM around 0, 226.077598874 Da}; Biotin-tyramide on Y; G-PTM-D modifications count = 1; precursor mass tolerance(s) = {±5.0000 PPM around 0, 361.146012788 Da} product mass tolerance = ±20.0000 PPM. max mods per peptide = 2; max modification isoforms = 1024; precursor mass tolerance = ±5.0000 PPM; product mass tolerance = ±20.0000 PPM; report PSM ambiguity = True. The combined search database contained 82427 non-decoy protein entries including 0 contaminant sequences. The following search settings were used: protease = trypsin or GluC, search for truncated proteins and proteolysis products = False; maximum missed cleavages = 2; minimum peptide length = 7; maximum peptide length = unspecified; initiator methionine behavior = Variable; fixed modifications = Carbamidomethyl on C, Carbamidomethyl on U; variable modifications = Oxidation on M; max mods per peptide = 2; max modification isoforms = 1024; precursor mass tolerance = ±5.0000 PPM; product mass tolerance = ±20.0000 PPM; report PSM ambiguity = True. The combined search database contained 82427 non-decoy protein entries including 0 contaminant sequences. The mass spectrometry proteomics data have been deposited to the ProteomeXchange Consortium via the PRIDE partner repository with the dataset identifier PXD068036 and reviewer access as follows: **Username:** and **password:** VkQAuYBtx8C7

## Acknowledgements

The authors acknowledge Dr. Jennifer Liddle (U. Conn Proteomics Core) for help with mass spectrometry instrumentation. The authors further acknowledge all Borniger lab members who provided valuable feedback on this manuscript.

## Notes

<u>COI Statement</u>: The authors report no conflicts of interest related to this work.

<u>Funding Sources</u>: J.C.B. is supported by an NIH R01CA286651, an AACR Breast Cancer Research Foundation – NextGen Grant for Transformative Cancer Research (20-20-26-BORN), and a Department of Defense Idea Development Award (W81XWH2210871). J.L.B. is supported by an NIH S10 High-End Instrumentation Award 1S10-OD028445-01A1, which supported this work by providing funds to acquire the Orbitrap Eclipse Tribrid mass spectrometer housed in the University of Connecticut Proteomics & Metabolomics Facility.

### Competing Interest Statement

The authors have declared no competing interest.

