## Supplementary Figure 1 for "APEX2 and TurboID Define Unique Subcellular Proteomes"

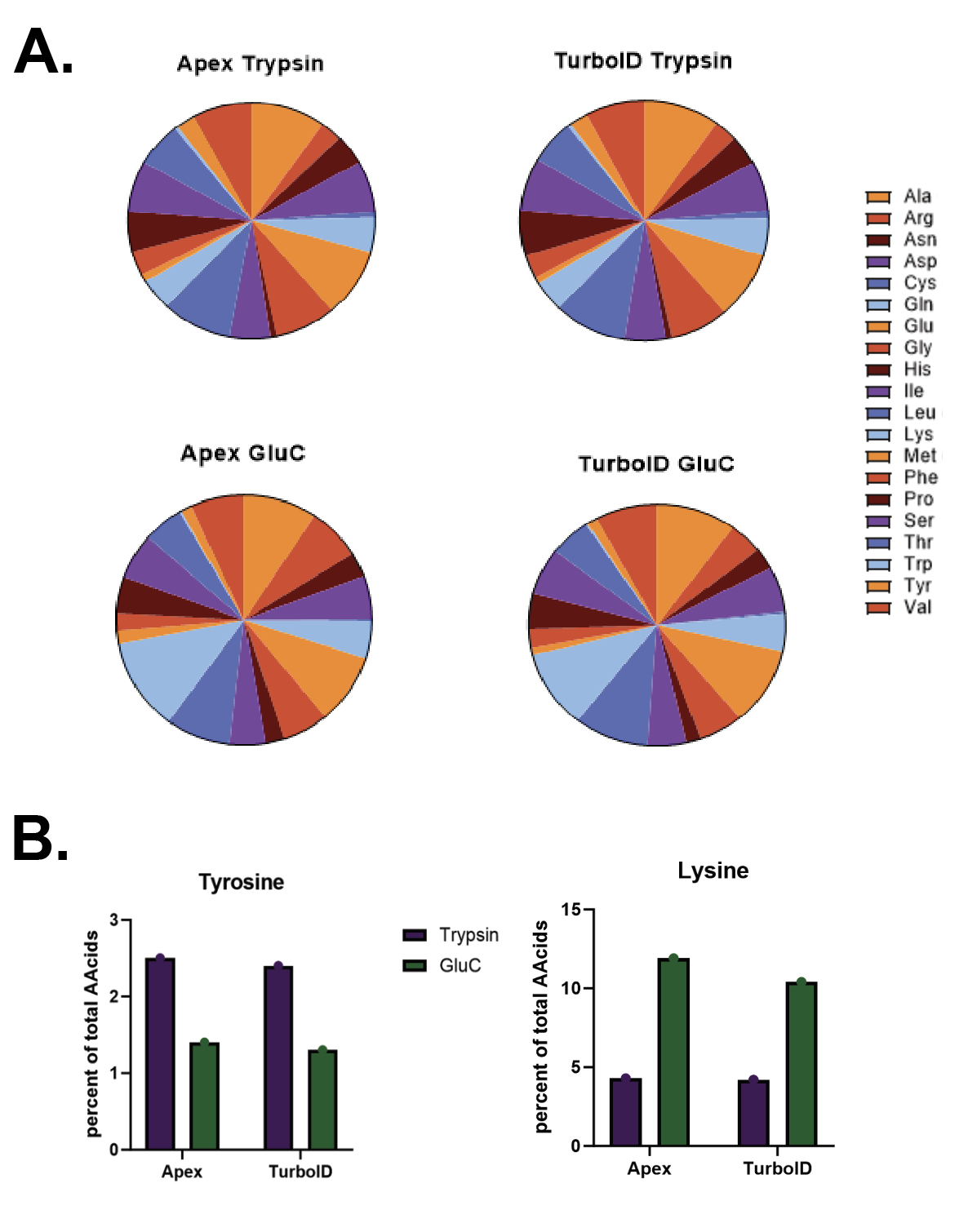

**Supplemental Figure 1: Frequency of amino acids.** A). Percentage of each amino acid detected across all peptides identified by mass spectrometry B). Percentage of total amino acids detected that are either tyrosine or lysine residues for each proximity labeling approach and each protease treatment.
